# Transcriptomic analysis with the progress of symbiosis in ‘crack-entry’ legume *Arachis hypogaea* highlights its contrast with ‘infection thread’ adapted legumes

**DOI:** 10.1101/251181

**Authors:** Kanchan Karmakar, Anindya Kundu, Ahsan Z Rizvi, Emeric Dubois, Dany Severac, Pierre Czernic, Fabienne Cartieaux, Maitrayee DasGupta

## Abstract

In root-nodule symbiosis, rhizobial invasion and nodule organogenesis is host controlled. In most legumes, rhizobia enter through infection-threads and nodule primordium in the cortex is induced from a distance. But in dalbergoid legumes like Arachis hypogaea, rhizobia directly invade cortical cells through epidermal cracks to generate the primordia. Herein we report the transcriptional dynamics with the progress of symbiosis in A. hypogaea at 1dpi: invasion; 4dpi: nodule primordia; 8dpi: spread of infection in nodule-like structure; 12dpi: immature nodules containing rod-shaped rhizobia; and 21dpi: mature nodules with spherical symbiosomes. Expression of putative orthologue of symbiotic genes in ‘crack-entry’ legume A. hypogaea was compared with infection thread adapted model legumes. The contrasting features were (i) higher expression of receptors like LYR3, EPR3 as compared to canonical NFRs (ii) late induction of transcription factors like NIN, NSP2 and constitutive high expression of ERF1, EIN2, bHLH476 and (iii) induction of divergent pathogenesis responsive PR-1 genes. Additionally, symbiotic orthologues of SymCRK, FLOT4, ROP6, RR9, NOOT and SEN1 were not detectable and microsynteny analysis indicated the absence of RPG and DNF2 homologues in diploid parental genomes of A. hypogaea. The implications are discussed and a molecular framework that guide ‘crack-entry’ symbiosis in A. hypogaea is proposed.

## INTRODUCTION

Nitrogen fixing root-nodule symbiosis (RNS) allows plants to house bacterial diazotrophs in an intracellular manner (Kistner and Parniske, 2002). The establishment of RNS involves rhizobial invasion in the root epidermis and nodule organogenesis in the root cortical cells. The most common invasion strategy is through root hair curling and infection thread (IT) formation where the nodule primordia are induced from a distance (Gage, 2004). Invasion through IT is widespread among temperate legumes like *Vicia sp., Trifolium sp.* and *Pisum sp*. Model legumes like *Lotus japonicus* and *Medicago truncatula* also undertake IT mediated rhizobial invasion (Geurts and Bisseling, 2002; Oldroyd and Downie, 2004, 2006). The alternate mode of rhizobial invasion is known as ‘crack-entry’ where the rhizobia enter through natural cracks at the lateral root base in an intercellular manner. Approximately 25% of all legume genera are adapted to ‘crack-entry’ and it is a characteristic feature of some subtropical legumes belonging to dalbergoid/genistoid clades like *Arachis sp., Aeschynomene sp.* and *Stylosanthes sp.*(Sprent and James, 2007). In these legumes, rhizobia directly access the cortical cells for development of their nodule primordia and the infected cells repeatedly divide to develop the mature nodule with aeschynomenoid infection zone (IZ) devoid of uninfected cells (Boogerd and Rossum, 1997; Fabre et al., 2015). Very limited information is available for ‘crack-entry’ legumes from dalbergioid/genistoid clade which are basal in their divergence within the papilionoideae.

Investigations on model legumes that are adapted to infection thread mediated rhizobial invasion have unravelled the molecular basis of RNS. The host responses are initiated by Nod-factor (NF) receptors *Lj*NFR1/*Mt*LYK3 and *Lj*NFR5/*Mt*NFP (Madsen et al., 2003; Radutoiu et al., 2003; Arrighi, 2006; Smit et al., 2007). Another NF induced receptor *Lj*EPR3 was shown to monitor rhizobial exopolysaccharide (EPS) in *L*. japonicus, indicating a two-stage mechanism involving sequential receptor-mediated recognition of NF and EPS signals to ensure host symbiont compatibility (Kawaharada et al., 2015). Downstream to NFRs is the ‘SYM pathway’ consisting of the receptor kinase *Lj*SYMRK/*Mt*DMI2 (Endre et al., 2002; Stracke et al., 2002), the predicted ion-channel proteins *Lj*Castor and *Lj*POLLUX/*Mt*DMI1(Ané et al., 2004; Imaizumi-Anraku et al., 2005), the nucleoporins *Lj*NUP85 and *Lj*NUP133(Kanamori et al., 2006; Saito et al., 2007), the Ca2+/calmodulin-dependent protein kinase *Lj*CCaMK/*Mt*DMI3 (Lévy et al., 2004; Tirichine et al., 2006), and the transcription factor *Lj*CYCLOPS/*Mt*IPD3(Messinese et al., 2007; Yano et al., 2008). Nodulation-specific transcription factors (TFs), such as *Mt*NSP1/*Lj*NSP1, *Mt*NSP2/*Lj*NSP2, *Mt*ERF1 and *Mt*NIN/*Lj*NIN function downstream of the ‘SYM pathway’ and are involved in transcriptional reprogramming for initiation of RNS (Schauser et al., 1999a; Kaló et al., 2005; Smit et al., 2005; Middleton et al., 2007).

Transcriptome analyses during symbiosis in several legumes like *M. truncatula, L. japonicus, Glycine max* and *Cicer arietinum* provided molecular insights into symbiosis that uses the ‘root hair’ mode entry (Takanashi et al., 2012; Limpens et al., 2014; Parida et al., 2016; Yuan et al., 2017a). These resources have been useful in determining gene expression associated with different stages of RNS, for example symbiont recognition, onset of infection, meristem activity for nodule organogenesis and functional differentiation of infected cells (Breakspear et al., 2014; Roux et al., 2014). The outcome in most cases highlighted numerous novel genes co-expressed with known markers for symbiosis. Nodule transcriptome in *M. truncatula* highlighted that transcription factor genes essential for the control of the root apical meristem were also expressed in the nodule meristem (Roux et al., 2014). Transcriptomic analysis of infected root hairs further revealed involvement of genes regulating cell cycle, flavonoid biosynthesis and auxin signaling during IT initiation (Breakspear et al., 2014). It also enabled identification of genes and processes previously undetected in whole-root studies or in forward genetic analyses. In *M. truncatula* transcriptomic analysis of symbiotic mutants further improved our understanding of gene expression associated with specific processes of symbiosis (Moreau et al., 2011; Jardinaud et al., 2016b). But unlike in model legumes, transcriptomic profiling for species that utilize ‘crack entry’ has rarely been reported to date and the symbiotic pathway of ‘crack entry’ nodulation is largely unknown.

An earlier report have listed several differentially expressed genes (DEGs) at an early stage of symbiosis in a nodulation competent *A. hypogaea* in comparison to an incompetent variety (Peng et al., 2017). Our manuscript undertakes a systematic effort to study the transcriptomic dynamics with the onset and advancement of symbiosis at 5 different nodule developmental stages in *A. hypogaea* using uninfected roots (UI) as a reference. Primarily we focused on symbiotic genes that were characterised in model legumes and could be broadly classified according to their functional involvement in bacterial recognition, early signalling, infection, nodule organogenesis, functional differentiation and nodule number regulation. Expression of putative orthologues of these symbiotic genes in ‘crack-entry’ legume *A. hypogaea* is compared and contrasted with the corresponding expression profiles in *M. truncatula* and *L. japonicus* that undertake IT mediated symbiosis. And the outcome of the analysis is summarised in a model that can be further utilised for understanding nodulation dynamics in basal legumes.

## RESULTS

### Progress of symbiosis in Arachis hypogaea

Within three weeks after infection with *Bradyrhizobium sp*. SEMIA 6144, A. *hypogaea* roots developed spherical functional nodules. We followed the progress of symbiosis in *A. hypogaea* for 21 days to identify the distinct stages of development by ultrastructure analysis. There are rosettes of root hairs in the junction of taproot and lateral root that are reported to be important for bacterial invasion in *A. hypogaea* (Boogerd and Rossum, 1997). Within 1 day post infection (1dpi) rhizobia was found to be adhered to these root hairs (Fig. 1A-C). Within 4dpi, bump like primordial structures were noted at the lateral-root bases (Fig. 1D). The longitudinal sections of these primordia revealed one or more centrally-located defined pockets of rhizobia infected cells that were surrounded by uninfected cells (Fig. 1E). These pockets of rhizobia infected cells were distinct by having reduced calcofluor-binding ability, indicating that they are thin-walled. The intracellularised rhizobia within the infection pockets were undifferentiated and rod-shaped (Fig. 1F). The infection pockets observed at 4dpi act as infection zone (IZ) founder cells and it is their uniform division and differentiation that give rise to the distinct aeschynomenoid type IZ in mature nodules. There has not been a single case where uninfected primordium was noted, which is in accordance with the proposition of infection preceding development of aeschynomenoid nodules (Fabre et al., 2015). By 8dpi there was visible nodule-like structure at the lateral-root base (Fig. 1G). Ultrastructure analysis revealed that by 8dpi, the compactness of the primordial structure with defined pockets of infected cells was lost and the IZ started growing by division of the infected cells (Fig. 1H-I). By 12dpi there were white spherical nodules (Fig. 1J). At this stage the tissue organization turned aeschynomenoid where there were no uninfected cells in the infection zone (IZ) and the endocytosed rhizobia remained undifferentiated and rod-shaped (Fig. 1K-L). At 21dpi the nodules were mature and functional where the rhizobia differentiated within the plant derived peribacteroid membranes to develop spherical symbiosomes (Fig. 1M-O).

**Figure 1:**
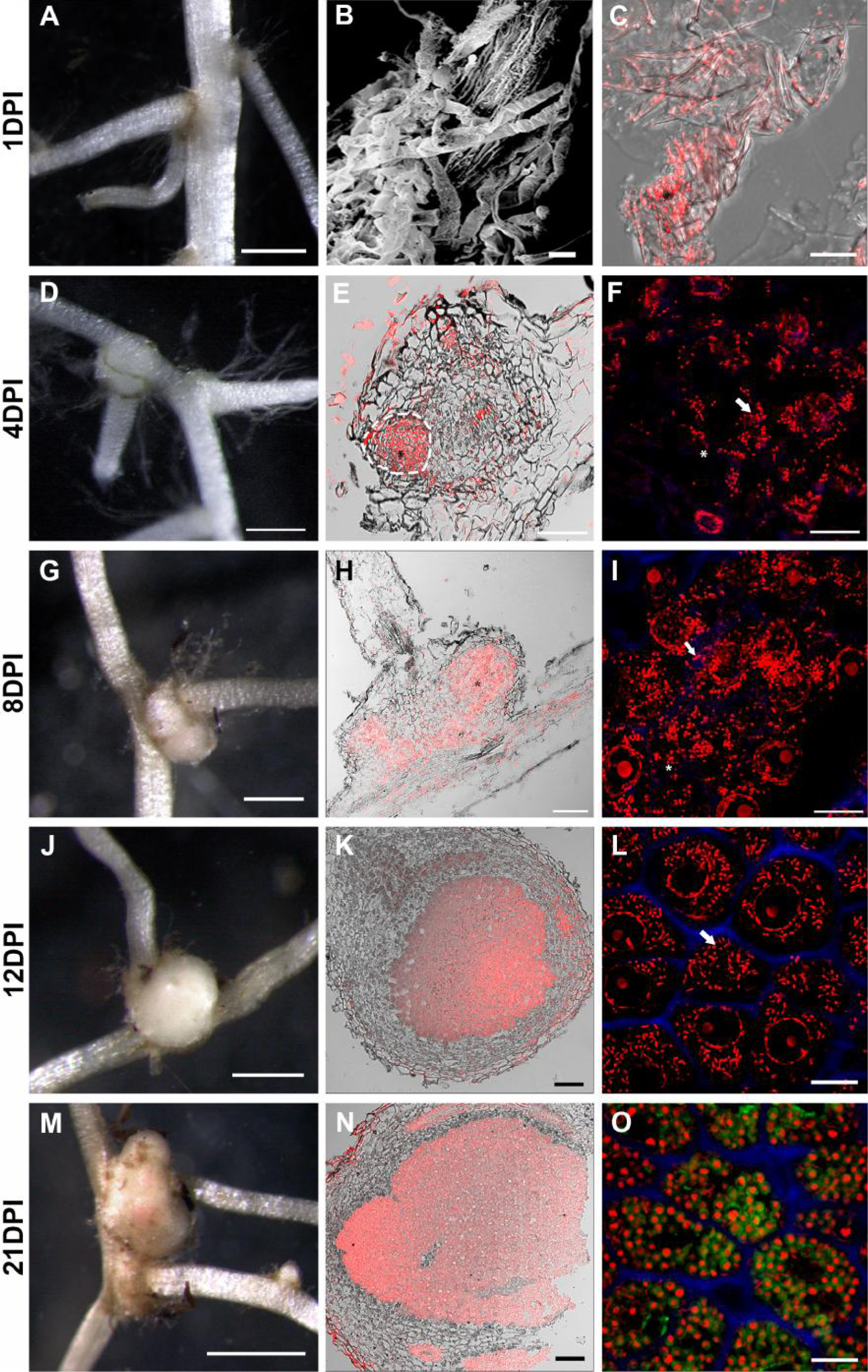
Progress of symbiosis in *Arachis hypogaea* infected with *Bradyrhizobium* sp. SEMIA6144. (A-C) Root harvested at 1dpi where (A) root junction, (B) SEM of inoculated root hair at lateral root junction and (C) CLSM image of lateral root junction. Section of nodule primordia at 4dpi (D-F) and 8dpi (G-I). Section of nodules at 12dpi (J-L) and 21dpi (M-O). Stereoimage (A, D, G, J and M); bright field and PI merged (C, E, H, K and N); PI + Calcofluor merged (F, I and L); PI + Calcofluor + syto9 merged (O). Dashed lines indicatethe infection zone in E and asterisk indicate identical position in E-F and H-I. Arrow indicates rod shaped rhizobia in F, I and L. Scale bar: 500μm (A, D, G and J), 1mm (M), 2μm (B), 100μm (E, H, K and N) and 10μm (C, F, I, L and O). PI (red), Calcofluor (blue) and Syto9 (green).

### Transcriptome analysis with the progress of symbiosis in *Arachis hypogaea*

Ultrastructural analysis revealed 5 distinct stages during the progress of symbiosis in *A. hypogaea* 1dpi: recognition and invasion; 4dpi: primordia formation; 8dpi: nodule-like structure; 12dpi: immature nodules with rod-shaped rhizobia; and 21dpi: mature nodules with spherical symbiosomes. To probe into the expression of genes associated with the progress of symbiosis, RNA was extracted from these stages. As a reference (0dpi) the uninfected (UI) roots were used. RNA-seq was done in triplicate for these six stages using Illumina single-end sequencing technology (Illumina Hiseq 2000 SR50). The genomic data from *Arachis duranensis* (AA) and *Arachis ipaensis* (BB) that are two wild diploid parents of *A. hypogaea* were used to assess the quality and coverage of the assembled transcriptomes. A total of 1,429,876,614 raw reads of 50bp (~71.5Gb) were generated with an average of 88,029,386 reads per library. This was 600 times the total size of transcript sequences (109.0 Mb) of *A. hypogaea* for both AA and BB genomes and gave an average coverage of 36 times per library. The proportion of clean reads among the total acquired reads was more than 91.34% (Table 1). The filtered reads were simultaneously mapped to the AA and BB genomes where the overall accepted mapping rate per library ranged from 80.15% to 89.98%, with an average mapping rate of 86.42% with *A. duranensis* (AA) and 86.65% with A. ipaensis (BB). For both AA and BB genome about 66% reads aligned to a gene exon in an unambiguous way, whereas the rest 33% reads aligned outside exon.

**Table 1:** Summary of Raw Illumina Sequencing and filtered reads after trimming and alignment of reads to AA (*Arachis duranensis*) and BB (*Arachis ipaensis*) genomes in each library

| Samples | Raw Reads | Filter reads (%) | % of Total Mapped Reads |  |
| --- | --- | --- | --- | --- |
|  |  |  | *Arachis duranensis*<br>(AA Genome) | *Arachis ipaensis*<br>(BB Genome) |
| UI | 57913214 | 95.08% | 82.96% | 86.71% |
|  | 65769652 | 94.88% | 87.78% | 84.34% |
|  | 85558982 | 90.59% | 85.29% | 86.19% |
| 1DPI | 61747256 | 95.00% | 88.80% | 87.51% |
|  | 62925897 | 91.38% | 88.78% | 89.98% |
|  | 74542718 | 91.68% | 87.85% | 88.76% |
| 4DPI | 96807458 | 89.21% | 88.46% | 84.42% |
|  | 97275402 | 90.46% | 86.61% | 87.71% |
|  | 79984160 | 91.21% | 85.24% | 86.25% |
| 8DPI | 89987084 | 90.09% | 86.03% | 87.19% |
|  | 11535441 | 89.85% | 86.20% | 83.03% |
|  | 77031650 | 91.50% | 87.81% | 88.95% |
| 12DPI | 57198145 | 90.28% | 86.77% | 88.08% |
|  | 74743668 | 92.33% | 88.47% | 89.71% |
|  | 96613689 | 89.38% | 86.12% | 87.20% |
| 21DPI | 65329428 | 90.03% | 87.81% | 86.81% |
|  | 100254251 | 90.71% | 84.45% | 85.42% |
|  | 70839550 | 90.46% | 80.15% | 81.44% |

The expression level of each assembled transcript sequence was measured through FPKM (Fragments per kilo-base per million reads) values. The DEGs in the 5 different stages of symbiosis were evaluated by the significance of differences in their expression with respect to UI roots using false discovery rate (FDR) < 0.05, P-value <0.05 and fold change |log2 ratio| ≥ 1 (Supplementary Table 1). Since a single UI was used for all the stages, there could be representations of some DEGs that are associated with normal root development within the experimental time frame. Comparison between upregulated and downregulated DEGs at different stages is shown in a Venn-Diagram in Fig. 2A. A total of 2745 genes were up-regulated (↑1297:AA,;↑1448:BB) and a total 20259 genes are down-regulated (↓9552:AA;↓10707:BB) with the progress of symbiosis in *A. hypogaea* of which 59 genes (↑29:AA;↑30:BB) were commonly upregulated and 2054 genes (↓1015:AA;↓1039:BB) were commonly downregulated in all the 5 stages of symbiosis. From the Venn-diagram we identified those genes that were first upregulated or downregulated at a particular stage though their subsequent regulations could be different. The number of such genes upregulated or downregulated at each stage from AA or BB genome is shown in Supplementary Table 2. Differentially expressed genes show clear transcriptional shifts at these stages and the diverse expression patterns of these genes are indicated in a heatmap (Fig. 2B). The major expression profiles are shown in expanded heatmaps and line graphs in Supplementary Fig. 1. Hierarchical clustering (Fig. 2B) and Principal Component Analysis (PCA) (Supplementary Fig. 1C) both suggested that there is substantial overlap between the transcriptome of infected roots of 1dpi and 4dpi where rhizobial invasion and primordia formation occurs. Similarly there was overlap between transcriptome of immature nodules at 8dpi and 12dpi where the primordia structurally develop into a nodule. Finally the transcriptome from mature nodules at 21dpi was distinct from all other stages where nodule matures to its functional form. These expression waves indicated that there could be 3 major transcriptional programs during the inception and maturation of symbiosis in *A. hypogaea*. The actual number of genes upregulated or downregulated from AA or BB genome at 1dpi was 337 and 8046 (↑156:↓3826:AA;↑181:↓4219:BB), at 4dpi was 388 and 4643 (↑180:↓2167:AA;↑208:↓2476:BB), at 8dpi was 641 and 1587 (↑297:↓735:AA;↑344:↓852:BB), at 12dpi was 502 and 1548 (↑240:↓751:AA;↑262:↓797:BB) and for mature nodules at 21dpi was 877 and 4435 (↑423:↓2141:AA;↑454:↓2294:BB). This analysis shows that within 1dpi-4dpi the number of downregulated DEGs were ~18-fold higher than the number of upregulated DEGs whereas in other stages the difference was ~3-5fold. Previous reports on symbiosis associated transcriptome of other legumes also indicated the prevalence of higher number of downregulated transcripts (Boscari et al., 2012; Yuan et al., 2017b). But the significantly high number of downregulated transcripts in early stages of nodule inception could be a unique feature for *A. hypogaea* RNS.

**Figure 2:**
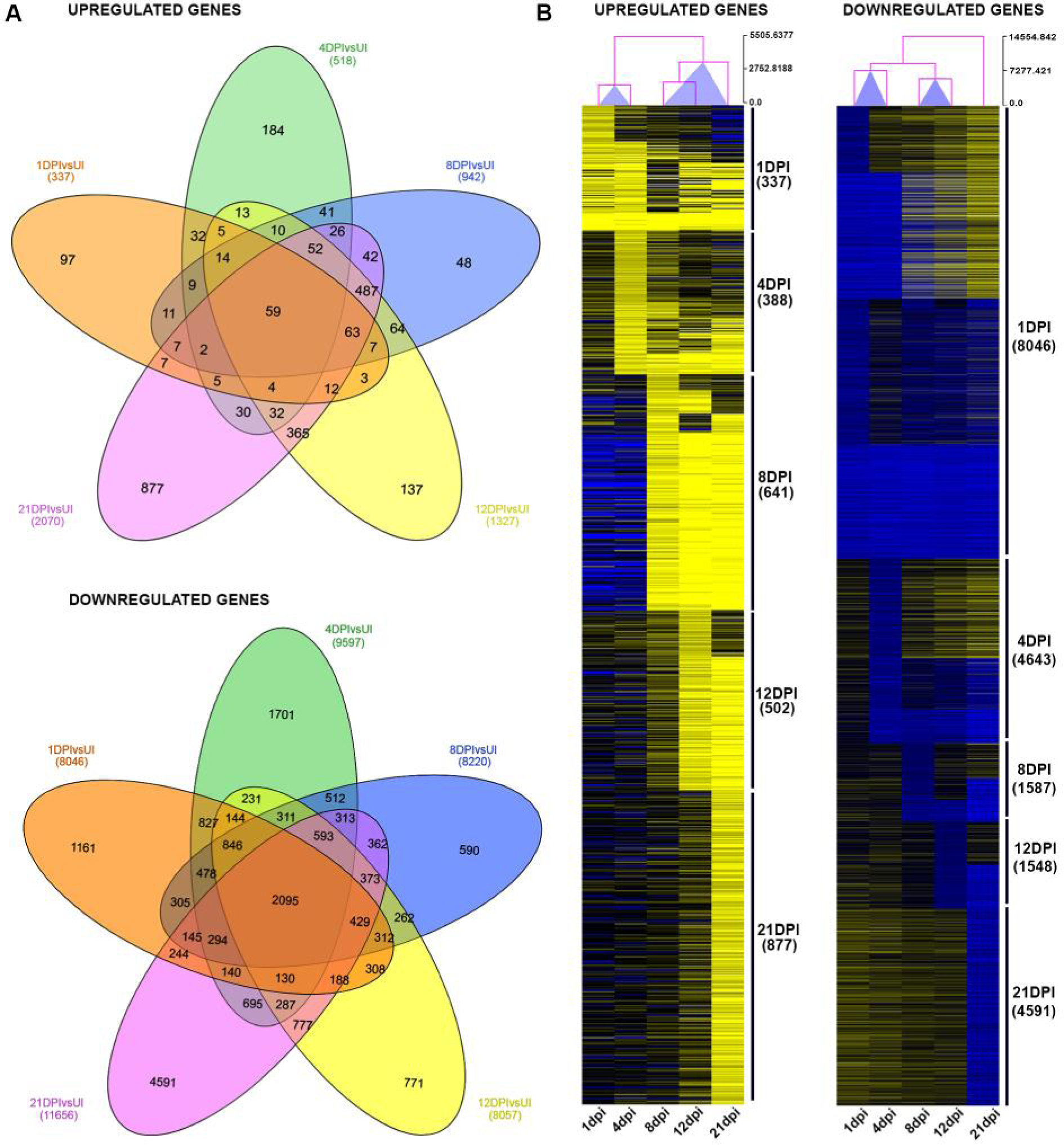
Comparison of differentially expressed genes (DEGs) at 5 different stages during the progress of symbiosis in *A. hypogaea*. The total DEGs at 1dpi, 4dpi, 8dpi, 12dpi and 21dpi was identified by EdgeR analysis from both AA and BB genomes. (A) Venn diagram showing comparison between upregulated and downregulated DEGs. (B) Heat map of the hierarchical cluster analysis of the same DEGs. Columns represent different time points and the arborescence above them indicates similarity. Scale bar represents log2 fold change in gene expression. Numbers beside the heat-map indicates the number of DEGs at different time point (see Supplementary Table 2).

### Functional analysis of DEGs

GO and KEGG terms that are significantly enriched in our DEGs are indicated in Supplementary Fig. 2. Among the 1248 enriched GO terms there was a major representation of defense response genes. 470 and 31 such defense related GO terms were enriched in downregulated and upregulated DEGs respectively (Supplementary Table 1). Accordingly, KEGG analysis of plant- pathogen interaction pathways show that most genes involved in pattern-triggered immunity (PTI) was notably down-regulated (Fig. 3A-B; Supplementary Table 3). The Flagellin sensing receptor (FLS2) mediated MAP kinase pathway (*MAPK*) however remained active along with a subset of Cyclic nucleotide-gated ion channels (*CNGCs*) and genes encoding Respiratory burst oxidase (*Rboh*) proteins. A subset of genes involved in the effector triggered immunity (ETI) also remained active during symbiosis, for example the genes encoding R proteins like *RPM1*, RPS2, *RPS5*, Pti1 kinase, and the pathway regulators like *SGT1, HSP90* and *EDS1* (Supplementary Table 3). Intriguingly there was a significant upregulation of genes encoding PR-1 proteins which are members of Cysteine-rich secretory proteins, Antigen 5, and Pathogenesis-related 1 proteins (CAP) superfamily (Breen et al., 2017). The PR-1 proteins upregulated during symbiosis clustered away from the PR-1 proteins that were reported to be upregulated in defence responses indicating the symbiosis associated PR-1 proteins to be divergent in nature (Fig. 3C). There are two PR-1 proteins that clustered with defence responsive PRs and these *PR-1* genes were not upregulated during symbiosis further confirming the symbiotic PR-1s to be distinct. PR-1 proteins harbour an embedded defence signalling peptide (CAP-derived peptides or CAPE) where CNYxPxGNxxxxxPY is considered as a functional motif that mark cleavage of these bioactive peptides (Breen et al., 2017). The cleavage site is conserved in both classes of PR-1 proteins suggesting the CAPE peptides could also be generated from the divergent PR-1 proteins synthesized during symbiosis (Fig. 3C-D, Supplementary Table 7). Since genes encoding CAP proteins are marker genes for the salicylic acid signaling pathway and systemic acquired resistance we also checked the SA/JA pathways to further understand the symbionts responsive signaling in A. hypogaea. As shown in Fig. 3A the JA pathway was completely downregulated but the SA responsive genes like *TGA1* and *NPR1* were up-regulated. Thus symbiotic *PR-1* gene expression could be justified by the activation of the SA mediated signaling.

**Figure 3:**
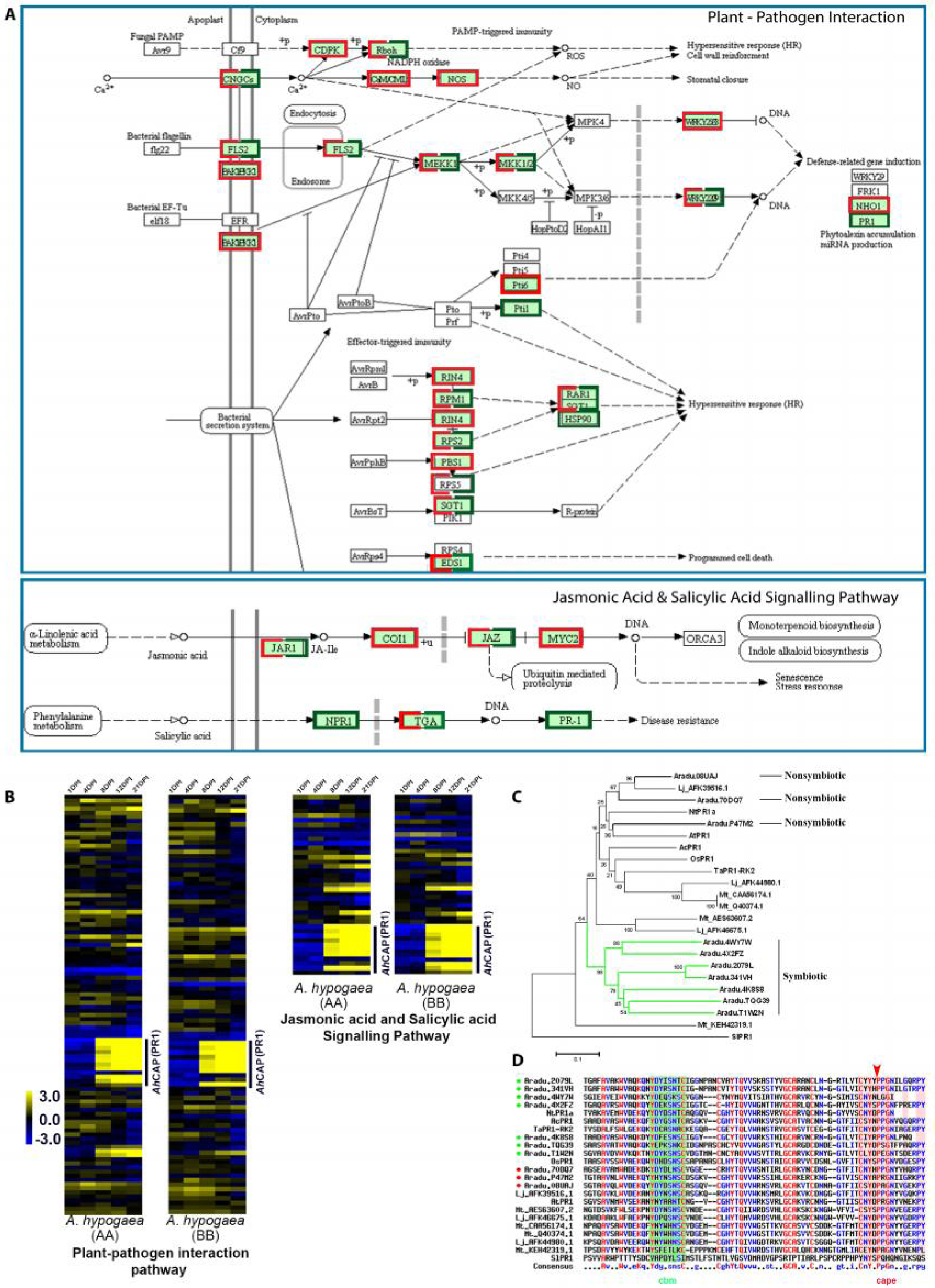
DEGs associated with plant-pathogen interaction and jasmonic acid-salicylic acid signaling pathways. (A) KEGG analysis of upregulated (green) and downregulated (red) DEGs in the respective pathways. (B) Heat-map of the DEGs that are annotated in KEGG pathway analysis. (C) Neighbour-joining phylogenetic tree of all the annotated CAP proteins using nontruncated amino acid sequences where green branch denotes divergent upregulated CAP-PR1 proteins. (D) CLUSTALW sequence alignment of CAPE peptides using Multalin where arrow indicates cleavage site for CAPE peptide; CBM and CAP domain are annotated by colored boxes and bullets indicates *A. hypogaea* CAP peptides (upregulated: green and downregulated: red).

Several genes are reported to be expressed in nodulating roots by comparing the transcriptome profiles of non-nodulating and nodulating lines of A. hypogaea (Peng et al., 2017). The list includes known symbiotic genes like *NIN* and *CLE13* and other genes like *NF-YA10*, *Myb127*, a receptor kinase, a soluble kinase, a F-BOX protein, transcription factors of SHI-family and a lectin (Supplementary Fig. 3; Supplementary Table 4). All these genes were represented in our upregulated transcriptome which thereby revalidates the importance of expression of these genes during the onset of symbiosis in *A. hypogaea*.

### Expression profiles of putative orthologues of symbiotic genes

Our final objective was to understand the expression of the putative orthologues of symbiotic genes in *A. hypogaea* that are characterized in the model legumes like *M. truncatula* and *L. japonicus*. A total of 71 such genes were chosen and functionally classified on the basis of their demonstrated role during nodulation in model legumes (Fig. 4; Supplementary Table 5). BLAT search on *A. ipaensis* and *A. duranensis* genome identified 63 genes that were annotated in PeanutBase and 6 genes that are yet to be annotated like *NSP2, ERN1, ERF1, VPY, ENOD40* and *RSD*. Homologues of *MtRPG* and *MtDNF2* could not be detected by BLAT search in either of the two parental genomes of *A. ipaensis* and *A. duranensis* which could be due to incomplete genome sequence or the quality of the gene model predictions. Microsynteny analysis in Phytozome indicated the absence of *MtRPG* orthologues in *A. ipaensis*, whereas in *A. duranensis* the entire geneblock corresponding to *MtRPG* was not detectable (Supplementary Fig. 4A). Griesmann et al., 2018 in a recent report also indicated the loss of *RPG* gene in *A. ipaensis* genome. Similar microsynteny analysis also revealed the absence of *MtDNF2* in *A. ipaensis* genome but indicated the presence of a singleton in *A. duranensis* (Supplementary Fig. 4B). Multiple alignment did not reveal any resemblance of this singleton to MtDNF2 suggesting its absence or fragmentation (Supplementary Fig. 4B). Moreover, we could not identify any homologous contigs corresponding to *MtRPG* and *MtDNF2* in our RNAseq data using Burrow- Wheller Aligner, thus confirming its absence in *Arachis hypogaea* transcriptome.

**Figure 4:**
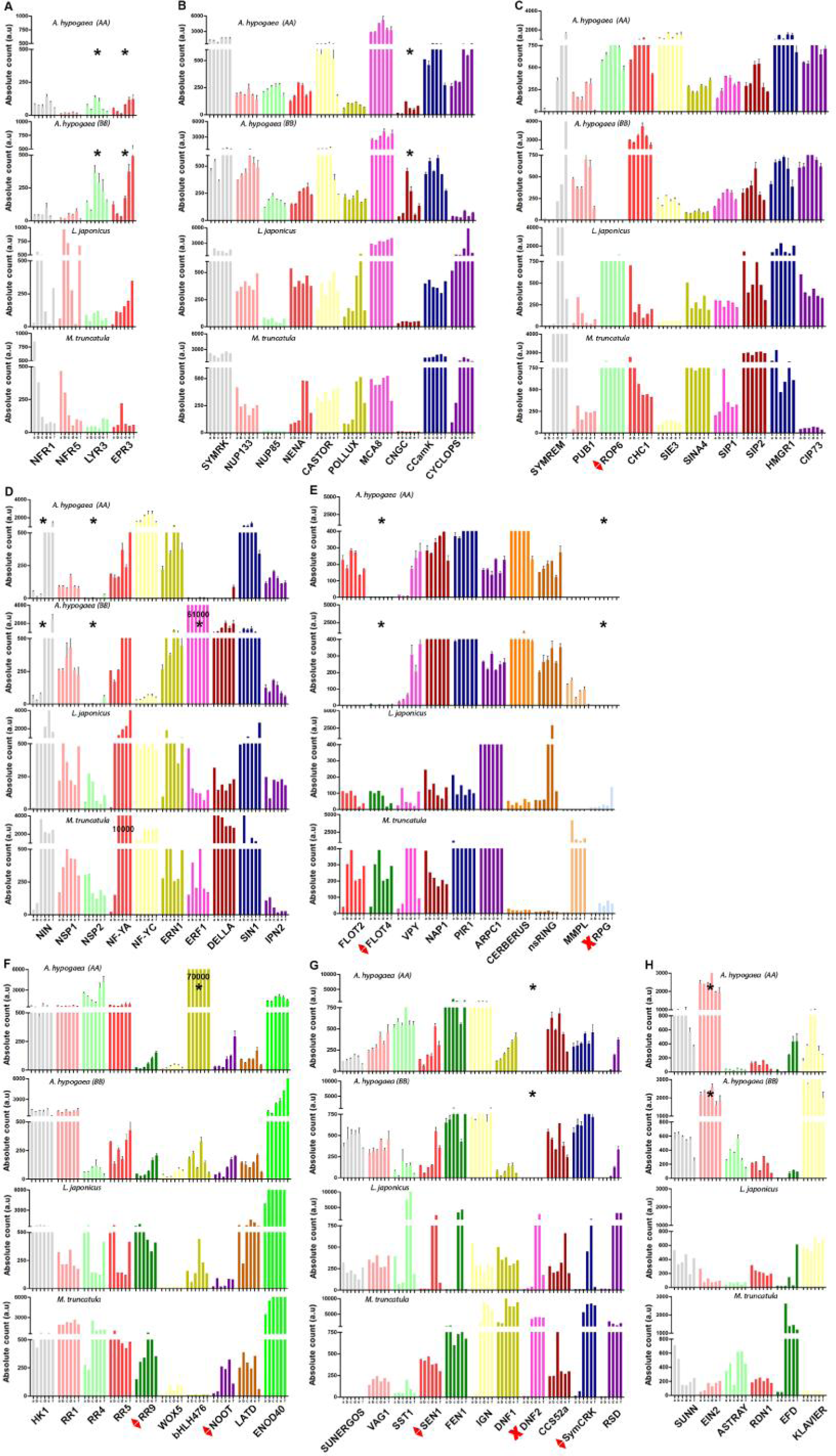
Comparative Expression Pattern of 71 Symbiotic genes in *A. hypogaea, M. truncatula* and *L. japonicus*. For *A. hypogaea* histogram represents normalized RNA-seq reads (FPKM) of symbiotic orthologous genes aligned to *A. duranensis* and *A. ipaensis* genomes. For *M. truncatula* and *L. japonicus* histogram represent microarray data retrieved from the respective Gene Expression Atlas Affymetrix database i.e MtGEA and LjGEA (Benedito et al., 2008; Verdier et al., 2013). Expression values of symbiotic genes (in arbitraty units, a.u.) during nodulation are grouped as (A) Bacterial recognition, (B) SYM pathway and early signaling, (C) Interactors, (D) Early transcription factors, (E) Infection, (F) Nodule organogenesis, (G) Nodule differentiation, (H) Nodule number regulation (dpi: days post inoculation by *Bradyrhizobium sp.* SEMIA6144 for *A. hypogaea, Sinorhizobium meliloti* for *M. truncatula* and *Mesorhizobium loti* strain R7A for *L. japonicus*). a,b,c,d,e and f in x-axis represented 0,1,4,8,12 and 21dpi respectively for *A. hypogaea*; 0,3,6,10,14 and 20dpi respectively for *M. truncatula*; 0,1,3.7,14 and 21dpi respectively for *L. japonicus*. Asterisk indicates genes that are differentially expressed in *A. hypogaea*. Red diamond: expression of non-symbiotic orthologue in *A. duranensis* and *A. ipaensis* and Red Cross: absent in diploid genome.

Orthology for the identified symbiotic genes were checked by reciprocal BLAST and sequence alignments. Symbiotic orthologues were identified for 63 out of 69 genes where the *A. hypogaea* homologues clustered with other legumes in the corresponding gene trees (Supplementary Fig. 5). Exceptions were the homologues for *MtSymCRK, MtFLOT4, LjROP6, MtRR9, MtNOOT* and *LjSEN1* where the *A. hypogaea* genes clustered with nonlegumes. For comparison of expression between the symbiotic or non-symbiotic orthologues in *Arachis sp*. with their counterpart in model legumes, we used the microarray data derived from the *M. truncatula* gene expression atlas (MtGEA) (http://mtgea.noble.org/v2/) (Benedito et al., 2008) and *L. japonicus* gene expression atlas (LjGEA) (https://ljgea.noble.org/v2/) (Verdier et al., 2013). For the symbiotic genes that are characterised from one of these model legumes, reciprocal BLAST was done to identify the orthologue in the other (Supplementary Table 5). The pattern of expression of these genes with the progress of symbiosis was compared between *A. hypogaea* and model legumes. Both absolute expression (Fig. 4) and relative (log2 fold) expression values (Supplementary Fig. 6) were analysed so that high and constitutively expressed genes were not ignored. The notable exceptions are mentioned below and the analysed data is summarised in a model in Fig. 5. The contrasting genes are highlighted along with the genes that are absent or divergent in sequence.

**Figure 5:**
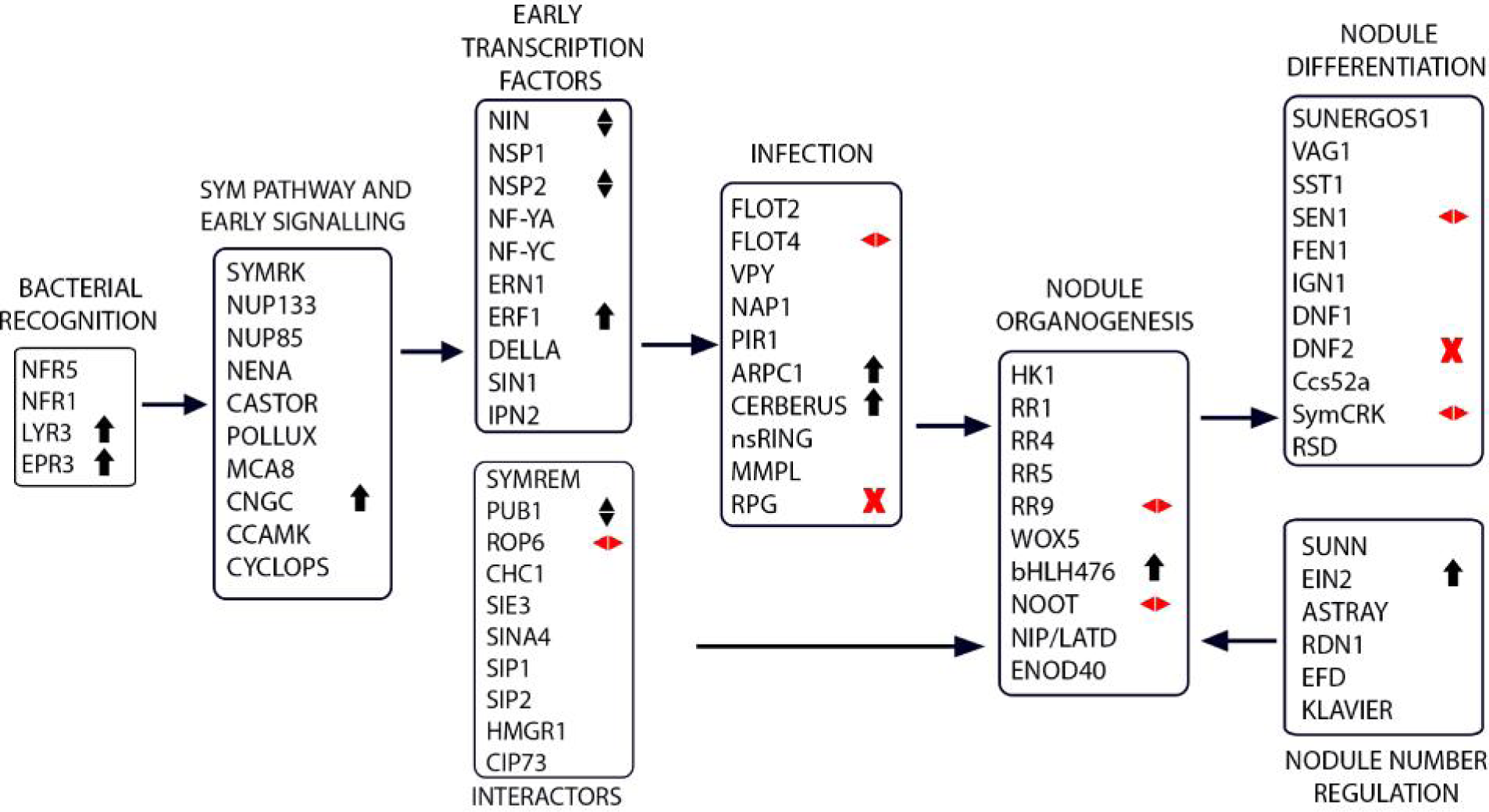
Simplified model for the symbiotic signaling pathway in *A. hypogaea*. Symbiotic genes identified in *M. truncatula* and *L. japonicus* are listed and classified according to their main symbiotic functions. The homologues of these genes in *A. hypogaea* that are distinct from model legumes are marked Red Cross: absent in diploid genome, Red diamond: symbiotic orthologue absent, Black up-head arrow: upregulation, Black diamond: divergent expression.

In the recognition module, expression of LCO-binding *AhLYR3* and exo-polysaccharide (EPS) binding *AhEPR3* were significantly high as compared to the orthologues of canonical Nod-factor LysM domain containing receptors *AhNFR1/5* (Fig. 4A, Fig. 5). Most of the SYM pathway members and their interactors and the early transcription factors showed similar trends in expression in both A. hypogaea and in model legumes for both inducible and constitutive genes (Fig. 4B-D). Exception was the gene encoding an orthologue of cyclic nucleotide-gated channel *MtCNGC* (Charpentier et al., 2016) which was significantly upregulated in *A. hypogaea* (Fig. 4B). Among the interactors, the orthologue of an E3 Ubiquitin ligase, *MtPUB1* was constitutively expressed in *A. hypogaea* whereas in the model legumes its expression is upregulated only upon bacterial invasion (Fig. 4C). In model legumes, TFs like *NIN* and *NSP2* are upregulated with bacterial invasion in early stages (Schauser et al., 1999b; Kaló et al., 2005). In contrast, upregulation of *AhNIN* (Singh et al., 2014)(Singh et al., 2014)(Singh et al., 2014) was noted at 8dpi onwards, and *AhNSP2* expression was induced only at 21dpi in mature nodules (Fig. 4D, Fig. 5). The expression of ethylene responsive TF, *AhERF1* appeared to be very high and constitutive with the progression of symbiosis. On the other hand, in model legumes the expression of *ERF1* was significantly reduced (*L. japonicus*) or upregulated (*M. truncatula*) upon bacterial invasion.

Among the genes associated with rhizobial invasion, the homologue for *MtRPG* (Arrighi et al., 2008), a factor responsible for rhizobium directed polar growth of ITs was absent in A. hypogaea (Supplementary Fig. 4A, Fig. 5). Additionally, symbiotic orthologues of *LjROP6*, a Rho-like GTPase that is responsible for IT elongation in cortex was not detected in *Arachis sp.*, though the expression of its nonsymbiotic homologue was significantly high during symbiosis (Fig. 4C, Supplementary Fig. 5). Among the flotillins that are required for lipid raft formation during infection (Haney and Long, 2009; Ke et al., 2012), symbiotic orthologues of *MtFLOT2* but not *MtFLOT4* could be detected (Fig. 4E, Supplementary Fig 5). Accordingly, *AhFLOT2* showed higher expression during symbiosis in comparison to *AhFLOT4*. The expression of the orthologue of *LjCERBERUS*, a WD-40 repeats U-box protein essential for the progress of infection in L. *japonicus* (Yano et al., 2009), was significantly high in A. hypogaea as compared to its expression pattern in model legumes (Fig. 4E).

Among the factors associated with nodule development, the homologues of *MtDNF2* encoding a phosphatidyl-inositol phospholipase C-like protein (Bourcy et al., 2013), is absent in *Arachis sp*. genome (Supplementary Fig.4B). In addition, symbiotic orthologues of a nonRD receptor kinase *MtSymCRK* (Berrabah et al., 2014), a type-A response regulator *MtRR9*, a metal transporter *LjSEN1* and a TF *MtNOOT* was also not detectable in *A*. duranensis and *A*. ipaensis genome (Fig. 4F-G, Supplementary Fig. 5). But significant expressions of the nonsymbiotic homologues of these genes were noted during the progress of symbiosis in *A. hypogaea*. On the other hand, the expression of orthologues for cytokinin inducible TF *MtbHLH476* (Ariel et al., 2012) and the ethylene responsive TF *MtEIN2* (Varma Penmetsa et al., 2008) (*sickle*) was found to be significantly higher in *A. hypogaea* as compared to their expression pattern in model legumes (Fig. 4F, 4H and Fig. 5). Apart from the indicated differences, relative expression patterns of all other symbiotic genes were found to be comparable between *A. hypogaea* and model legumes during progress of symbiosis indicating their conserved roles.

### qRT-PCR validation of the RNAseq results

To validate the RNAseq results we did qRT-PCR analysis using *AhActin* as the reference gene. The stability of expression of AhActin during symbiosis in *A. hypogaea* was reported earlier (Morgante et al., 2011; Kundu and DasGupta, 2017). The expression was evaluated for 10 symbiotic genes from different modules that are described in Fig. 5. Additionally expression was evaluated for 5 more genes that were described to be important for *A. hypogaea* symbiosis by Peng et al., 2017. The results were in agreement with the transcriptional profile data for 83 out of 90 (92%) data points (Supplementary Fig. 7). The fold-change of expression did not always match between RNAseq and qRT-PCR, but the trend of expression was similar in most of the time points for all the genes. No change of expression was noted for any of these genes in uninoculated roots corresponding to different infection time points, further confirming the expression of these genes to be specifically associated with symbiosis.

### PCA of symbiotic gene expression in Arachis hypogaea and model legumes

PCA was done to check if there was a signature in the expression pattern of symbiotic genes in ‘crack-entry’ legume *A. hypogaea* that contrasts with the model legumes where rhizobial entry is IT mediated. Fig. 6 is a projection of differential expression of symbiotic genes from *A. hypogaea, M. truncatula* and *L. japonicus* for all the infection time points into first two principal components. Altogether, expression of around 87% genes were accommodated along dimension1 (PC1) and dimension 2 (PC2) in the analysis. Expressions of symbiotic genes that are constitutive or show minimal change are clustered near the origin for both *Arachis sp*. and the model legumes. These include the SYM pathway and most early signaling genes that are likely to be regulated at the post-transcriptional level. The genes that were noted to have a contrasting trend in expression between *A. hypogaea* and the model legumes in Fig. 4 were all placed in opposing or adjacent quadrants in the PCA. For example, *AhCNGC, AhNIN, AhNSP2, AhPUB1*, and *AhCERBERUS* were clustered separately from their counterparts in model legumes. On the other hand there are some factors whose contrasting trends in expression between *A. hypogaea* and model legumes was not obvious in Fig. 4, but PCA highlighted their differences. For example *AhVPY, AhENOD40, AhWOX5, AhRR1* and *AhRR4* were placed in opposing or adjacent quadrants from their counterparts in model legumes. These factors highlighted by PCA can also be considered to be differentially adapted for ‘crack-entry’ mediated rhizobial invasion and aeschynomenoid nodule development in *A. hypogaea.*

**Figure 6:**
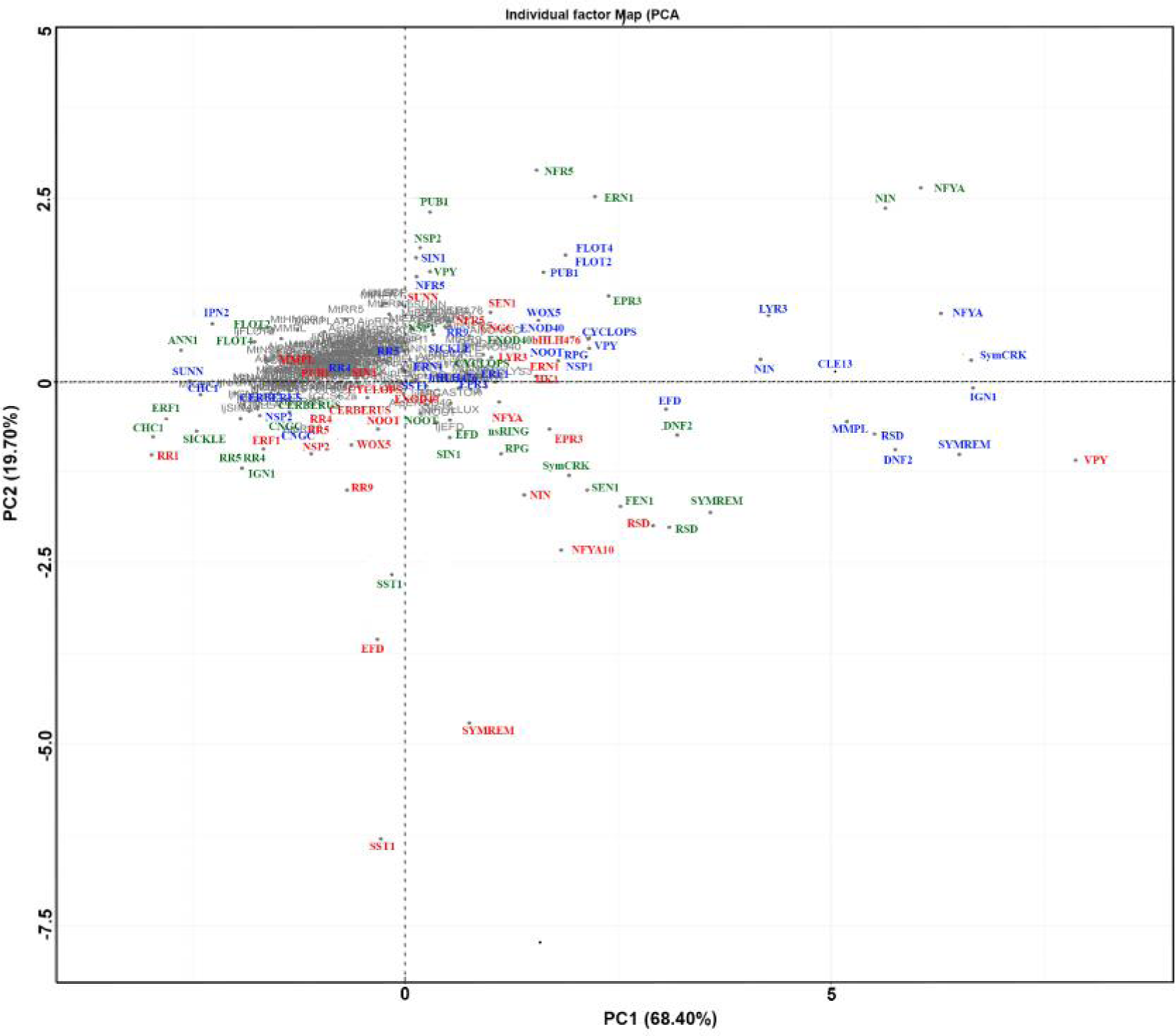
Principal Component Analysis (PCA) of differentially expressed symbiotic genes. Principal components of the variability in expression values for the symbiotic genes at all the infection time points in *A. ipaensis* (red), *M. truncatula* (blue) and *L. japonicus* (green) was calculated using R-Package. The two principal components and their fraction of the overall variability of the data (%) are shown on the x-axis (PC1 68.40%) and the y-axis (PC2 19.70%). All the genes are highlighted by a single acronym and the details of the symbiotic genes are mentioned in Supplementary Table 5.

## DISCUSSION

This is the first systematic effort towards transcriptome profiling with the progress of symbiosis in a ‘crack-entry’ legume *A. hypogaea*. We could correlate the hierarchical shift in upregulated DEGs to the morphological changes occurring during different time points of nodule development and the process appeared to be governed by three major transcriptional programs. The first program is associated with rhizobial recognition and generation of nodule primordia by 1 −4dpi, the 2^nd^ program is associated with structural development of nodules by 8-12dpi and the 3^rd^ program is associated with functional maturation of nodules at 21dpi (Fig. 1–2, Supplementary Fig. 1, Supplementary Table 1). The comparison between expression patterns of putative orthologues of symbiotic genes with progress of symbiosis in *A. hypogaea* with model legumes highlighted the genes that are important or dispensable for the ‘crack-entry’ mediated root nodule symbiosis (Fig. 4–6, Supplementary Table 5). Additionally absence of symbiotic orthologues of certain symbiotic genes in *A. hypogaea* was highlighted (Supplementary Fig. 4-5). The probable implications of these distinguishing features in *A. hypogaea* are discussed in the light of their importance during ‘crack-entry’ nodulation.

The most significant observation in *A. hypogaea* symbiotic transcriptome was the significantly high expression of a group of genes encoding a divergent form of cysteine rich PR-1 proteins (Fig. 3). Upregulation of *AhCNGC* and *AhERF1* in the symbiotic transcriptome (Fig. 4) could explain the expression of *PR1* genes, as their role in SA induced PR gene expression is well documented (Solano et al., 1998; Asamizu et al., 2008; Moeder et al., 2011). PR-1 proteins are ubiquitous across plant species and are among the most abundantly produced proteins in plants in response to pathogen attack (Breen et al., 2017). As a result it is used as a marker for salicylic acid-mediated disease resistance in plants. Although differential expression of defense response genes belonging to GO:0006952 (defense related) and *PR-1*/*PR*-10 protein families has previously been reported for *M. truncatula* RNS (Jardinaud et al., 2016a), this is the first report where a divergent group of PR-1 proteins is shown to be associated with nodule development (Fig. 3, Supplementary Table 3 and 7). PR-1 proteins harbor a caveolin binding motif (CBM) that binds sterol and the embedded Pro-rich C-terminal peptide (CAPE) is involved in plant immune signaling (Breen et al., 2017). All the symbiotic PR-1s in *A. hypogaea* have both these conserved features but whether these CAPE peptides are actually derived from PR-1 proteins during symbiosis remains to be understood. It may be noted that in the IRLC clade of legumes as well as in a dalbergioid legume *Aeschynomene evenia* another group of cysteine rich peptides belonging to NCR family is responsible for bacterial differentiation associated with morphological change of symbionts (Van de Velde et al., 2010). Intriguingly, morphological change of symbionts from rod to spherical shape is also noted in *A. hypogaea* (Kundu and DasGupta, 2017). Similar to NCRs, whether the antimicrobial CAPE peptides are enrolled as symbiosis effectors in *A. hypogaea* remains to be understood.

Bacterial recognition is Nod-factor dependent in *A. hypogaea* and Nod-factor independent for *A. evenia* though rhizobial invasion in both these legumes are mediated through epidermal cracks (CHANDLER et al., 1982; Giraud et al., 2007; Ibáñez and Fabra, 2011). In accordance, the canonical Nod-factor receptor NFR1 was absent in A. evenia (Chaintreuil et al., 2016), but expressed in *A. hypogaea* (Fig. 4A, Fig. 5). Intriguingly, the expression of *AhLYR3*, orthologue of LCO binding LysM receptor *MtLYR3* (Fliegmann et al., 2016) was higher in A. hypogaea as compared to canonical *NFR1/5*, which is unlike model legumes where the expression of canonical *NFRs* were much higher than orthologues of *MtLYR3* (Fig. 4A). Since *LjLYS12*, an orthologue of MtLYR3 is important for recognition of pathogens (Kelly et al., 2017), it is tempting to suggest a possible involvement of *Ah*LYR3 in symbiont recognition in *A. hypogaea*. PUB1, a substrate of *LjNFR1* has a role in determining partner specificity (Mbengue et al., 2010). The constitutive expression of *AhPUB1* in *A. hypogaea* suggests that it might have such a role during the choice of symbionts (Fig. 4C). Another step in bacterial recognition is through perception and recognition of surface molecules present on the bacterial cell wall like exopolysaccharides (EPS), lipopolysaccharides (LPS), cyclic glucans and capsular polysaccharides (Via et al., 2016). EPS binding LysM receptor, *Mt*EPR3 has a role in bacterial recognition followed by IT elongation (Kawaharada et al., 2015) and its orthologue in *A. hypogaea* showed significant expression with progress of symbiosis (Fig. 4A). This receptor is absent in *A. evenia* transcriptome (Chaintreuil et al., 2016), indicating that it may not be important during crack invasion but rather it might have a role in symbiont recognition during *A. hypogaea* nodulation. Thus similar to infection thread adapted model legumes, expression of LysM receptors are also important for symbiont recognition during intercellular invasion of rhizobia in a ‘crack-entry’ legume like *A. hypogaea*.

Most of the genes that are important for IT progression in model legumes are significantly expressed during the progress of symbiosis in ‘crack-entry’ legume *A. hypogaea* (Fig. 4E). Exception is *Mt*RPG that has a role in polar growth of ITs in *M. truncatula* (Arrighi et al., 2008; Murray et al., 2011). Our results indicated absence of RPG homologue in *Arachis sp*. (Supplementary Fig. 4A), which is in accordance with a recent report where loss of RPG gene is indicated in *A. ipaensis* genome (Griesmann et al., 2018). Absence of *RPG* among the non- nodulators and ‘crack-entry’ legumes like *Arachis sp*. and *A. evenia* indicates that nodulation has evolved even at the loss of RPG gene (Chaintreuil et al., 2016). Along with *RPG*, symbiotic orthologue of *Mt*FLOT4 which is important for IT initiation and elongation (Haney and Long, 2009), was absent in both *A. evenia* (Chaintreuil et al., 2016) and divergent in *A. hypogaea* (Supplementary Fig. 5). Other factors that are important for IT progression in model legumes are *MtFLOT2* (Haney and Long, 2009), *MtVPY* (Murray et al., 2011), *LjCERBERUS* (Yano et al., 2009) and *LjARP1* (Hossain et al., 2012). Symbiotic orthologues for all these genes are present in *A. hypogaea* and they demonstrated similar expression pattern during symbiosis in comparison to the IT adapted model legumes indicating the importance of their additional roles during ‘crack- entry’ nodulation (Fig. 4). For example, *LjCERBERUS* is functionally associated with nodule organogenesis and *Lj*ARPC1 with cytoskeletal rearrangement during bacterial endocytosis (Yano et al., 2009; Hossain et al., 2012) and these additional functions of CEREBERUS and ARPC1 could be conserved in ‘crack-entry’ legumes.

Though cytokinin signaling is important for nodule development in both ‘crack-entry’ and IT adapted legumes (Gonzalez-Rizzo et al., 2006; Fabre et al., 2015; Kundu and DasGupta, 2017), PCA revealed distinct expression pattern for several downstream factors in *A. hypogaea* as compared to model legumes. For example, homologue of *MtRR1*, a type-B RR has a distinct expression pattern in *A hypogaea* and unlike model legumes, its downstream transcription factor *MtbHLH476* (Ariel et al., 2012) was highly expressed with the progress of infection in *A. hypogaea* (Fig. 4F, Fig. 6). The symbiotic orthologue of *MtRR9* (den Camp et al., 2011), a Type- A RR is absent in *A. hypogaea*, but the expression pattern of its nonsymbiotic homologue indicate its possible recruitment during symbiosis (Supplementary Fig. 5, Fig. 4F and Fig. 5). Apart from the RRs, the cytokinin responsive TFs like, *Lj*NIN (Tirichine et al., 2007), *Mt*NSP2 (Plet et al., 2011) that are central regulators for both nodule organogenesis and rhizobial infection do not show early expression during nodulation in *A. hypogaea* as compared to model legumes. Thus unlike in model legumes, the first transcriptional program that are associated with rhizobial recognition and generation of nodule primordia in *A. hypogaea* may not involve *NIN* and *NSP2*. Thus, cytokinin signaling pathway and its effectors appeared to be differentially adapted for regulating nodule development in ‘crack-entry’ legumes.

Several contrasting features were also noted among the symbiotic genes involved in nodule differentiation. For example, homologues for *MtDNF2* and symbiotic orthologues of *MtSymCRK* was not detectable in both *A. hypogaea* (Fig. 4G, Fig. 5, Supplementary Fig. 4B and 5) and *A. evenia* transcriptome (Chaintreuil et al., 2016). As both these factors are required for suppressing defense responses during bacteroid differentiation (Bourcy et al., 2013; Berrabah et al., 2014), the absence of *DNF2* and the divergence of *SymCRK* in ‘crack-entry’ legumes further indicate that the developmental program for aeschynomenoid legumes is distinct from the IT adapted legumes. *LjSUNERGOS* and *LjVAG1* codes for topoisomerases and are required for endoreduplication of plant cells (Suzaki et al., 2014; Yoon et al., 2014). Although the orthologues of *LjSUNERGOS* and *LjVAG1* were missing in *A. evenia* transcriptome, they are expressed in *A. hypogaea* highlighting the differences in nodule differentiation program between the Nod dependent (Fig 4G) and independent ‘crack-entry’ legumes (Chaintreuil et al., 2016).

Previous report on *A. hypogaea* transcriptomics (Peng et al., 2017), as well as our transcriptome (Supplementary Fig. 3, Supplementary Table 4) highlighted the upregulation of several AP2- domain containing ethylene responsive TFs during nodulation. *LjERF1* downregulates the expression of defense gene *LjPR-10* during symbiosis (Asamizu et al., 2008), whereas in *A*. hypogaea high expression of *AhERF1* is associated with high expression of a divergent group of PR-1genes indicating the differences in ethylene signaling between the IT adapted legumes and *A. hypogaea* (Fig. 4D). Also the expression of *AhEIN2*, the master regulator of ethylene signaling was found to be constitutively high in *A. hypogaea* at all post infection time points (Fig. 4H, Fig. 5). This predominance of ethylene signaling during *A. hypogaea* nodulation is in consistence with the previous observations in *Sesbania rostrata* where under submerged conditions ethylene signaling inhibited intracellular infection via infection threads while promoting intercellular infection via ‘crack-entry’. (Guinel and Geil, 2002; D’Haeze et al., 2003).

In summary, the comparative transcriptomic analysis have shed light on the contrasting expression profiles of DEGs in ‘crack-entry’ vs IT adapted symbiosis. Genes like *VPY, FLOT, CERBERUS* and *ARPC1* that are primarily known for their role in IT elongation are significantly expressed in *A. hypogaea* making it imperative to investigate their role in ‘crack-entry’ adapted legumes.

Similarly, the non symbiotic homologue of genes like *SymCRK, ROP6, RR9, NOOT* and *SEN1* are significantly expressed during symbiosis in *A. hypogaea* and it is important to know whether these genes were co-opted for symbiosis. Further investigations are also required to understand how cytokinin and ethylene signaling may have been differentially adapted in ‘crack-entry’ legumes. Finally, in absence of early induction of *NIN* and *NSP2* it is important to identify the factors that co-ordinates the rhizobial invasion and nodule primordia formation in ‘crack-entry’ legumes. The highlighted molecular intricacies can be further investigated for better understanding of ‘crack- entry’ nodule development. But overall, our investigations have revealed a considerable overlap in expression profiles of most of the symbiotic genes between a ‘crack-entry’ legume *A. hypogaea* and the infection thread adapted model legumes suggesting a functional conservation of most of the factors that govern the process of nitrogen fixing symbiosis irrespective of the mode of rhizobial entry in their host plants.

## MATERIALS AND METHODS

### Plant Materials and Sample Preparation

Five different developmental stages of *A.hypogaea* total infected roots, primordia/nodules and uninfected roots were used in this study (UI/0dpi, 1dpi, 4dpi, 8dpi, 12dpi and 21dpi). *A. hypogaea* JL24 strain seeds (from ICRISAT, INDIA) were surface sterilized with sodium hypochlorite solution (3% active chlorine) containing 0.01% Tween 20 for 15 min and rinsed five times with sterile deionized water and the sterile seeds were soaked in sterile deionized water for germination. Germinated seeds were then transferred in pots containing sterile vermiculite and soilrite at 25°C growth room for 7 days before inoculation with *Bradyrhizobium sp*. SEMIA 6144 (Sinharoy et al., 2009). *Bradyrhizobium sp*. SEMIA 6144 obtained from MIRCEN, Brazil and gifted by Adriana Fabra, Universidad Nacional de Rio cuarto, Cordoba, Argentina. Samples are harvested, cleaned and freezed in liquid nitrogen. Frozen samples are stored at −80°C for RNA isolation.

### Phenotypic analysis and microscopy

Images of whole-mount nodulated roots were captured using a Leica stereo fluorescence microscope M205FA equipped with a Leica DFC310FX digital camera (Leica Microsystems). Detached nodules were embedded in Shandon cryomatrix (Thermo scientific) and sliced into 30-μm thick sections with a rotary cryomicrotome CM1850 (Leica Microsystems). For confocal microscopy, sample preparation was done according to Haynes and associates (Haynes et al., 2004). Sections were stained with Calcofluor (Life Technologies), Propidium Iodide (Life Technologies) and Syto9 (Life Technologies). Images were acquired with a Leica TCS SP5 II AOBS confocal laser scanning microscope (Leica Microsystems). For confocal and scanning electron microscopy, sample preparation was done according to (Kundu and DasGupta, 2017). All digital micrographs were processed using Adobe Photoshop CS5.

### Isolation of total RNA

A total 100mg of frozen plant root was ground in liquid nitrogen, and total RNA was isolated using Trizol reagent (Invitrogen, USA). RNA degradation and contamination was detected on 1% agarose gels. RNA concentration was then measured using NanoDrop spectrophotometer (ThermoScientific). Additionally, RNA integrity was assessed using the Bioanalyzer 2100 system (Agilent Technologies, Santa Clara, CA, USA). Finally, the samples with RNA integrity number (RIN) values above 8 were used for library construction.

### Library construction and Sequencing

18 RNA library was prepared using an IlluminaTruSeq stranded mRNA sample preparation kit by MGX-Montpellier GenomiX core facility (MGX) France (https://www.mgx.cnrs.fr/). The protocol first requires the selection of polyadenylated RNAs on oligodT magnetic beads. Selected RNAs are chemically fragmented and the first strand cDNA is synthesized in the presence of actinomycin D. The second strand cDNA synthesis is incorporating with dUTP in place of dTTP which quenches it to the second strand during amplification. A 3’ ends adenylation was used to prevent fragments from ligating to one another during the adapter ligation process. The quantitative and qualitative validation of the library was performed by qPCR, ROCHE Light Cycler 480 and cluster generation and primary hybridization are performed in the cBot with an Illumina cluster generation kit. The sample libraries were sequenced on an IlluminaHiSeq 2000, sequencing by synthesis (SBS) technique performed by MGX, France and 50bp single-end reads for each library were generated(Fuller, 1995).

### Illumina Reads Mapping and Assembly

Quality control and assessment of raw Illumina reads in FASTQ format were done by FastQC software (Version 0.11.5) to obtain per base quality, GC content and sequence length distribution. Clean reads were obtained by removing the low quality reads, adapters, poly-N containing reads by using Trimmomatic v0.36 software (Bolger et al., 2014). Clean Reads are simultaneously aligned to the two wild peanut diploid ancestors *A*. duranensis (AA) and *A*. ipanensis (BB) reference genome by using TopHat2 version 2.0.13 which is a fast splice junction mapper for RNA-Seq reads where upto 3 mismatches were authorized per reads (Trapnell et al., 2010; Bertioli et al., 2015). It aligns RNA-Seq reads using the ultra high-throughput short read aligner Bowtie2 version 2.2.3, and then analyzes the mapping results to identify splice junctions between exons (Langmead et al., 2009). The alignment files were combined and analyzed into Trinity for genome-guided assembly (Grabherr et al., 2011).

The reference based assembly was compared to its respective transcript files from annotated reference genomes by using BLAT (Kent, 2002). An e-value cutoff of ‘1e^−05^’ was used to determine a hit. The annotated hits were furthermore analyzed in this study. Genome annotation files in generic feature format (GFF) are downloaded from peanut database (https://peanutbase.org/download) (Dash et al., 2016). Estimation of gene expression level of each annotated transcript was performed by StringTie v1.3.3 which takes sorted sequence alignment map (SAM) or binary (BAM) file for each sample along with genome annotation files (Pertea et al., 2015). Counting reads in unannotated genes is performed with HTseq-count after converting the coordinates of these genes (chromosomal positions) to a ‘GFF’ file. As the data is from a strand-specific assay, the read was mapped to the opposite strand of the gene. Resulted gene transfer format (GTF), normalized gene locus expression level as fragments per kilobase million (FPKM), transcripts per million (TPM), and count files for each sample were further analyzed for fold change analysis in gene expression levels. RNAseq reads alignments have been performed using Burrow-Wheller Aligner (BWA version 0.7.15-r1140) (Li and Durbin, 2009) against *MtRPG* and *MtDNF2* transcripts resulting in 100% read to be declared as unmapped.

### Identification of DEGs and functional Gene Ontology and KEGG pathway analyses of the DEGs

Before statistical analysis, genes with less than 2 values lower than one count per million (cpm) were filtered out. EdgeR 3.6.7 package was used to identify the differentially expressed genes (Robinson et al., 2010). Data were normalized using “Trimmed mean of M-values (TMM)” method. Genes with adjusted p-value less than 5% (according to the FDR method using Benjamini-Hochberg correction) and|log2 (fold change)|>1 was called differentially expressed. Venn-diagram are generated using (http://www.interactivenn.net/) (Heberle et al., 2015) and hierarchical heatmap is generated usingTM4MeV (http://mev.tm4.org and http://www.tigr.org/software/tm4/mev.html) (Howe et al., 2011) the values from the venn diagram (Supplementary Table 2). PCA was performed by a web-tool ClustVis (https://biit.cs.ut.ee/clustvis/) (Metsalu and Vilo, 2015). Principal components of the variability of expression data was calculated using PCA Method R-Package. Detailed functional annotation and explanations of DEGs were extracted from gene ontology database (http://www.geneontology.org/) (Ashburner et al., 2000) and GO functional classification analysis was done using software WEGO (http://wego.genomics.org.cn/cgi-bin/wego/index.pl) (Ye et al., 2006). The GO terms for DEGs in genome annotation were also retrieved from the ‘GFF’ file downloaded at PeanutBase website (http://peanutbase.org). To identify important and enriched pathways involved by the DEGs, the DEGs were assigned to the Kyoto Encyclopedia of Genes and Genomes (KEGG) pathways using the web server (http://www.genome.jp/kaas-bin/kaas_main) (Kanehisa and Goto, 2000) against A. duranensis and A. ipaensis gene datasets. Enriched KO and GO terms are obtained by a developed Python script which uses hypergeometric test and Bonferroni corrected p-Value < 0.05.

### Identification of symbiotic orthologous gene in *A. hypogaea*

Candidate symbiotic genes were identified in *A. duranensis, A. ipaensis, L. japonicus* and *M. truncatula* using BLASTN searches with reported nucleotide sequence of genes from L. japonicus and *M*. truncatula. The homologous genes were searched in *A. duranensis* and *A. ipaensis* at PeanutBase (http://peanutbase.org), *M. truncatula* Mt4.0v1 genome was searched at *M. truncatula* gene expression atlas (MtGEA) (http://mtgea.noble.org/v2/) and the *L. japonicus* v3.0 genome was searched at *L. japonicus* gene expression atlas (LjGEA) (https://ljgea.noble.org/v2/). Initial searches were conducted with E-value = e^−5^. The results were manually validated for the presence of an orthologous gene in an open reading frame and searched for orthologues using BLASTP. Orthology of the genes were validated by generating neighbor joining phylogenetic tree using amino acid sequences and nucleotide sequences (ENOD40) in MEGA 6.0 (Tamura et al., 2013). Syntenic analysis of RPG and DNF2 between the legumes was performed using Legume Information System (LIS) Genome context viewer (https://legumeinfo.org/lis_context_viewer/index.html#/search/lis/).

### qRT PCR validation

Total RNA (500 ng) was reverse-transcribed by using Super-ScriptIII RT (Life Technologies) and oligo (dT). RNA quantity from each sample in each biological replicate was standardized prior to first-strand cDNA synthesis. qRT-PCR was performed by using Power SYBR Green PCR Master Mix (Applied Biosystems) using primers as designed using software Oligoanalyser (IntergratedDNA Technology) (Supplementary Table 6). Calculations were done using the ΔΔcycle threshold method using *AhActin* as the endogenous control. The reaction were run in Applied biosystems 7500 Fast HT platform using protocol: 1 cycle at 50°C for 2 mins, 1 cycle at 95°C for 5 min, 40 cycles at 95°C for 30 sec, 54°C for 30 sec, 72°C for 30 sec followed by melt curve analysis at 1 cycle at 95°C for 1 min, 55°C for 30 sec, and 95°C for 30 sec. A negative control without cDNA template was checked for each primer combination which was designed using OligoAnalyzer 3.1 (https://www.idtdna.com/calc/analyzer). Results were expressed as means standard error (SE) of the number of experiments.

### Data Availability

The raw FASTQ files for the 18 libraries were deposited in the Gene expression omnibus (GEO) of NCBI under accession number GSE98997.

## Author’s Contribution

Project planning: A.K. and M.D.G. Sample preparation: K.K. and A.K.; Microscopy: A.K.; Preparation of RNA: K.K. Production of Illumina libraries, sequencing and transcriptome assembly: E.D, D.S.; Analysis of transcriptome: K.K. and A.Z.; Analysis of symbiotic transcriptome: K.K and A.K.; Critical analysis of data: P.C and F.C. Writing of the manuscript: A.K., K.K. and M.D.G. All authors approved the manuscript.

## Acknowledgement

This work was funded by Grants from Govt. of India: IFCPAR/CEFIPRA (IFC/5103-4/2014/543); DBT-CEIB (Centre of Excellence and Innovation in Biotechnology, BT/01/CEIB/09/VI/10); DBT-IPLS (BT/PR14552/INF/22/123/2010; fellowship to K.K and A.Z.R (IFCPAR/CEFIPRA: IFC/5103-4/2014/543); fellowship to A.K (Council of Scientific and Industrial Research, CSIR-09/028[0756]/2009-EMR-I).

